# Landscape structure is a key driver of protist diversity along elevation gradients in the Swiss Alps

**DOI:** 10.1101/2022.04.13.488160

**Authors:** Christophe V.W. Seppey, Enrique Lara, Olivier Broennimann, Antoine Guisan, Lucie Malard, David Singer, Erika Yashiro, Bertrand Fournier

## Abstract

**Context:** Human-induced changes in landscape structure are among the main causes of biodiversity loss. Despite their important contribution to biodiversity and ecosystem functioning, microbes - and particularly protists - remain spatially understudied. Soil microbiota are most often driven by local soil properties, but the influence of the surrounding landscape is rarely assessed.

**Objectives:** We assessed the effect of landscape structure on soil protist alpha and beta diversity in meadows in the western Swiss Alps.

**Methods:** We sampled 178 plots along an elevation gradient representing a broad range of environmental conditions and land-use. We measured landscape structure around each plot at 5 successive spatial scales (i.e. neighbourhood windows of increasing radius, ranging from 100 to 2000 m around a plot). We investigated the changes of protist alpha and beta diversity as a function of landscape structure, local environmental conditions and geographic distance.

**Results:** Landscape structures played a key role for protist alpha and beta diversity. The percentage of meadows, forests, or open habitats had the highest influence among all landscape metrics. The importance of landscape structure was comparable to that of environmental conditions and spatial variables, and increased with the size of the neighbourhood window considered.

**Conclusions:** Our results suggest that dispersal from neighbouring habitats is a key driver of protist alpha and beta diversity which highlight the importance of landscape-scale assembly mechanisms for microbial diversity. Landscape structure emerges as a key driver of microbial communities which has profound implications for our understanding of the consequences of land-use change on soil microbial communities and their associated functions.

## 1 Introduction

With about 75% of the global land area being impacted by human activities (Ellis and Ramankutty, 2008), land-use change is one of the main causes of terrestrial biodiversity loss (Brondizio et al., 2019; Cardinale et al., 2012; Mace et al., 2018; Sala et al., 2000). Land-use changes have modified landscape structure through destruction, degradation, and/or fragmentation of natural habitats with negative consequences on local biodiversity and the associated ecosystem functions and services (Brondizio et al., 2019). Anthropogenic modifications of landscape structure are likely to continue in the future to meet the increasing need of a growing human population (Mace et al., 2018; Tilman et al., 2011). In this context, understanding the impact of changes in landscape structure on ecosystems is required to maintain landscape biodiversity and services.

Microbes represent the dominant part of biodiversity on Earth (Locey and Lennon, 2016). They are also central actors to key ecosystem functions and services including carbon sequestration (Shakoor et al., 2020), litter decomposition (Jackson et al., 2017), nutrient turnover (Wagg et al., 2014), food production (Nosheen et al., 2021), water quality supply (Sagova-Mareckova et al., 2021) or regulation of biological populations (Kibblewhite et al., 2008). However, despite their important contribution to biodiversity and ecosystem functions, microbes are rarely considered in spatial studies assessing the impact of environmental conditions on biodiversity (e.g. Malard et al., 2022; Mod et al., 2020, 2021; Seppey et al., 2020), and landscape structure is considered even less frequently (Mennicken et al., 2020; Mony et al., 2020). As a result, microbial assembly processes and, especially, the importance of landscape structure for local microbial diversity and functionality remains unclear. This is a major knowledge gap in microbial ecology as well as in our understanding of the consequences of land-use changes.

Studies analysing the drivers of microbial diversity have primarily focused on local processes assuming that microbial community composition results from the effect of the local abiotic environment and from biotic interactions (Louca et al., 2018; Malard et al., 2022; Mod et al., 2020; Yashiro et al., 2018). These studies have shown the importance of environmental conditions such as temperature (Fierer et al., 2006; Mod et al., 2020), pH (Chu et al., 2010; Yashiro et al., 2016), and soil moisture (Sofi et al., 2016). Other studies have focused on biogeographical processes revealing patterns of microbial diversity (reviewed by Martiny et al., 2006), such as continental divide (Glaeser and Overmann, 2004; Louca, 2021), latitudinal gradient (Schwalbach and Fuhrman, 2005), or distance-decay relationships (Hanson et al., 2019; Schwob et al., 2021). However, these studies have mostly ignored the impact of regional-scale processes on local community composition (D’Amen et al., 2017; Leibold et al., 2004). Because landscape structure modulates the strength and type of impact of regional processes on local communities (Fournier et al., 2017, 2020), modern landscape ecology approaches (e.g. Riitters, 2019) have the potential to shed new light on the processes driving microbial community assembly, and particularly on the importance of dispersal from a regional pool of potential colonists.

Despite their potential, landscape ecology approaches applied to microbial communities remain scarce (Mony et al., 2020, 2022), mostly due to the difficulty to observe these organisms. Progress in DNA-based characterization of community composition now makes it easier to conduct landscape-scale surveys of microbial diversity (Kibblewhite et al., 2008). However, the few landscape ecology studies focusing on microbes are taxonomically and functionally biased towards parasitic organisms and pathogens, mostly fungi or bacteria (Cruz-Paredes et al., 2021; Mony et al., 2020). This line of research demonstrated the importance of landscape structures such as roads for the dispersal and distribution of pathogens (e.g. Bousset et al., 2018; Holdenrieder et al., 2004; Laine and Hanski, 2006). Yet, to our knowledge, the effect of landscape structure on microbial α and β diversity at large taxonomic scale (here, protists) has never been assessed. Because of the key functional importance of soil protists (Geisen, 2016; Seppey et al., 2017; Xiong et al., 2018), this gap limits our understanding of the consequences of land-use change on ecosystem structure and function.

The present study evaluates the effect of landscape structure on local soil protist α and β diversity across 178 non-forested open vegetation plots (mostly meadows) in the western Swiss Alps. We assess the effect of descriptors of landscape composition (i.e. the relative proportion of land cover types in the landscape, regardless of spatial distribution) and configuration (i.e. the spatial arrangement of land cover types in the landscape) on local microbial α and β diversity and compare it to environmental and spatial descriptors. We hypothesise that the structure of the landscape around a focal plot influences α and β protist diversity highlighting the importance of regional processes for local community assembly. We also expect that protist community dissimilarity (β diversity) correlates positively with geographic distance (distance-decay relationship) and that landscape, environmental and spatial descriptors contribute to this correlation. In addition, we examine the importance of scale for landscape effects on protist diversity by explicitly analysing the effect of landscape descriptors calculated at 5 successive spatial scales (neighbourhood window from 100 to 2000 m around a plot). If our analyses confirm our hypothesis, it would suggest that landscape-scale processes are key to understanding microbial community assembly and biodiversity patterns, as observed in larger-sized organisms (Betts et al., 2019; Martin et al., 2019).

## 2 Material & Methods

### 2.1 Study site and sampling design

We focus primarily on alpine meadows in the western Swiss Alps (Figure 1). As in most Western European mountain systems, meadows are threatened by both urbanisation and agricultural abandonment allowing forests to take over. As a result, the area occupied by meadows is decreasing and remains are increasingly fragmented (Dengler et al., 2014). These changes in landscape structure impact plants (Dengler et al., 2014) and, expectedly, protist diversity, and are likely to increase in the future (Pellissier et al., 2013). For these reasons, alpine meadows in the study area constitute a useful model ecosystem to study the impact of landscape structure on soil microbial communities.

**Figure 1:**
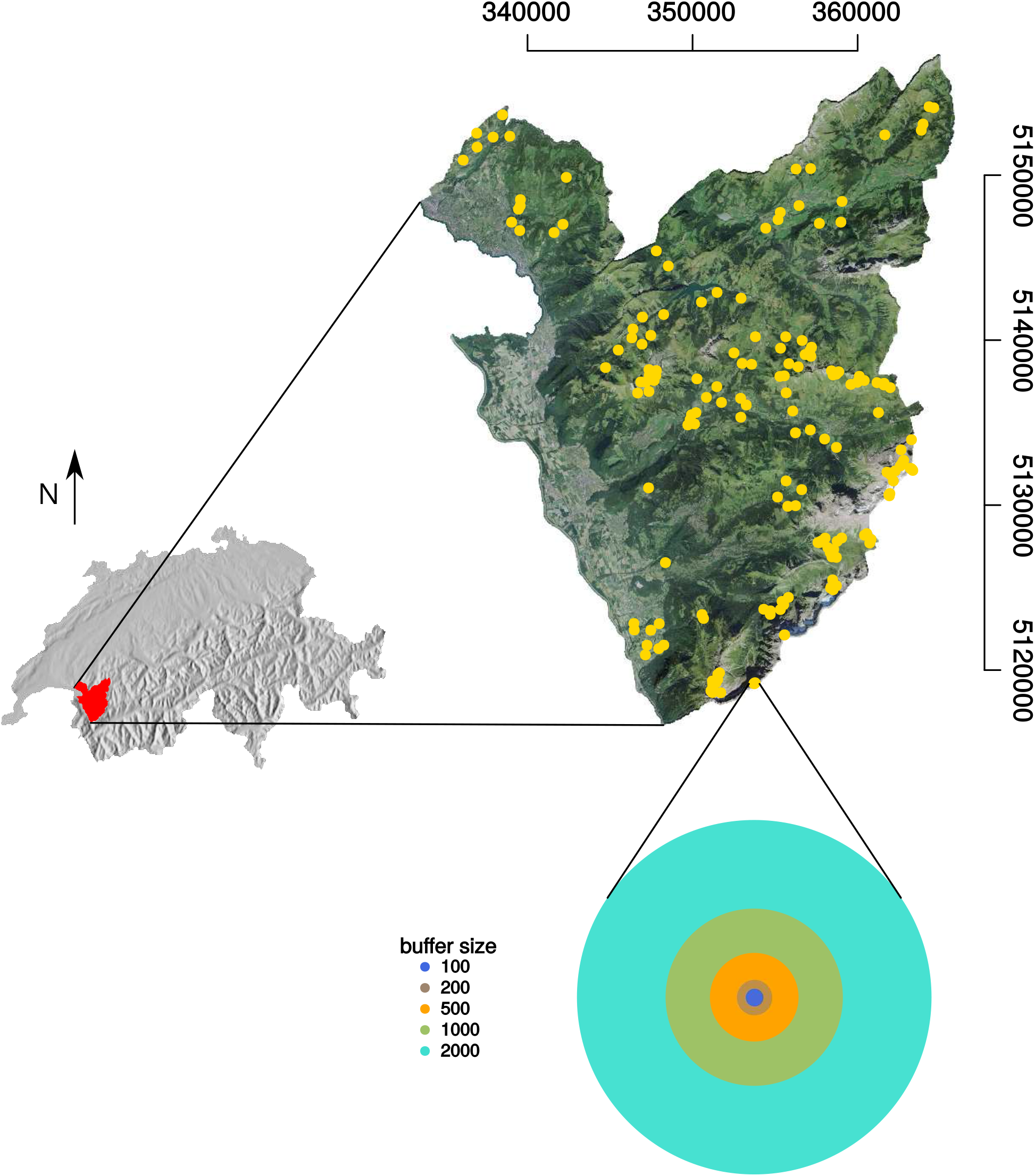
Map of the western Swiss Alps study area with the 178 selected plots’ locations (yellow dots). The lower panel shows the selected neighbourhood windows around each plot (100, 200, 500, 1000, 2000 m) used to calculate the different landscape metrics (see Table 1).

A total of 178 non-forest open vegetation plots (each of 2 m * 2 m) were sampled in the study area from July 4^th^ to September 1^st^ 2013 according to a sampling design stratified by altitude. For each sample, five soil cores of 100 g each soil cores were taken (from the four corners and centre of the plot, respectively, i.e. one ‘sample’). The soil cores were then mixed and kept below 4 °C until further processing. For more details, see Buri et al. (2020) and Yashiro et al. (2016).

### 2.2 Environmental descriptors

For each sample, 33 edaphic descriptors were measured (Table S1). Relative humidity (rh) and soil organic carbon (LOI) have been measured gravimetrically. Percentages of sand, silt, and shale were measured by laser granulometry. Electroconductivity (EC) and pH were measured in a soil and Milli-Q water slurry. Total phosphorus was measured by colorimetry after decarbonization. C/N ratio was calculated from total carbon and nitrogen content measured from spectroscopy (Carlo Erba CNS2500 CHN elemental analyzer). Total organic carbon (TOC) and mineral carbon (MINC) were measured by pyrolysis. The mineralogy was determined by X-Ray powder diffraction and fluorescence. A description of these analyses can be found in Buri et al. (2020) and Yashiro et al. (2016).

Topo-climatic descriptors were derived from data of the Swiss Meteorological Network (resolution 25 m; average trends for the period 1961-1990; https://www.meteoswiss.admin.ch) and Swisstopo (resolution 25 m https://www.swisstopo.admin.ch/). The descriptors were elevation (mnt25), slope steepness (slope), slope orientation (asp25) and the sinus of the orientation (aspvar), topographic position (topo), topographic wetness index (twi25), site water balance (swb), shortwave radiation per month (sumradyy), monthly mean precipitation sum (precyy), number of precipitation days per growing season (pday), monthly moisture index (mmind68), monthly average temperature (taveyy), annual degree-day above 0 ^°^C (ddeg0), and number of frost days during the growing season (sfroy) (Buri et al., 2020; Mod et al., 2020, 2021; Pradervand et al., 2014; Yashiro et al., 2016, 2018). In addition to these descriptors, bioclimatic descriptors were added from the Chelsa database (Karger et al., 2017) (Table S1).

For each environmental descriptor, the few missing values (1 rh, 4 spectroscopy, 3 mineralogy, 6 C/N when nitrogen content was below the detection limit) were inferred by nearest neighbour averaging (function `preProcess`, method ‘knnImpute’, package `caret` v. 6.0-86 Kuhn, 2020). The dimensionality of the environmental descriptor dataset was reduced using a principal component analysis. In order to capture 80% of the total variance in each dataset, we retained the first eleven PCA axes (Figure S1).

### 2.3 Landscape descriptors

The landscape descriptors have been calculated based on the 10 m resolution land cover raster derived from Sentinel-2 imagery (Malinowski et al., 2020) (http://s2glc.cbk.waw.pl/). This raster includes 13 land cover categories such as coniferous forest (representing 27.2% of the study area), herbaceous vegetation (25.6%), moors and heathlands (16.2%), and broadleaf tree forest (11.0%). A total of 13 descriptors commonly used in landscape ecology studies and representing different aspects of landscape configuration and composition were computed (see Table 1 for details). To explore the spatial scale at which landscape structure impacts soil protists diversity, each descriptor was calculated in various circular areas of different radius around the sampling coordinate (neighbourhood window size = 100, 200, 500, 1000, and 2000 m; Figure S2).

**Table 1:**
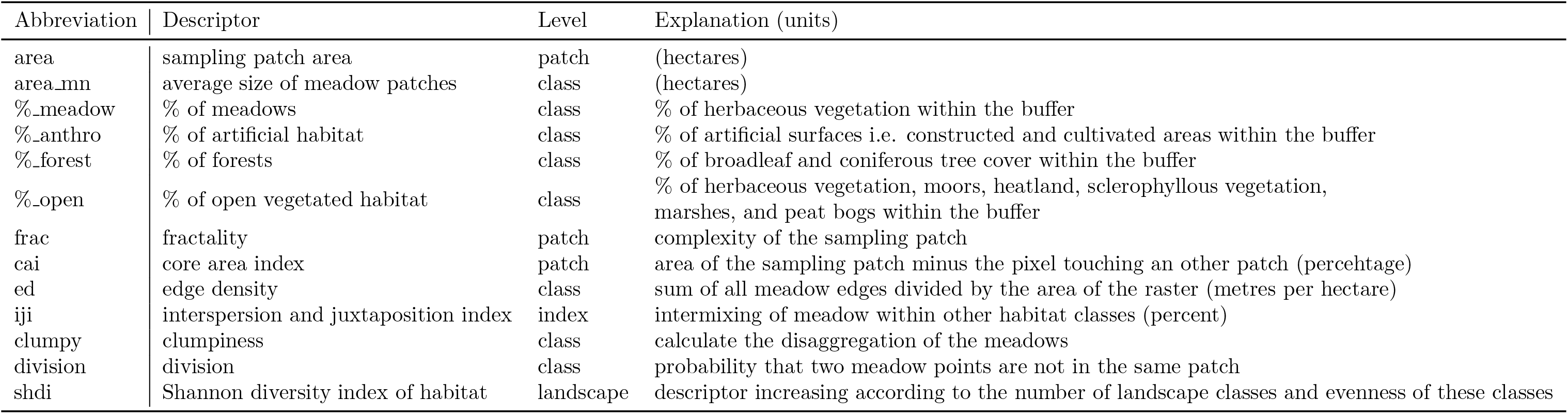
List of the selected landscape descriptors.

| Abbreviation | Descriptor | Level | Explanation (units) |
| --- | --- | --- | --- |
| area | sampling patch area | patch | (hectares) |
| area_mn | average size of meadow patches | class | (hectares) |
| %_meadow | % of meadows | class | % of herbaceous vegetation within the buffer |
| %_anthro | % of artificial habitat | class | % of artificial surfaces i.e. constructed and cultivated areas within the buffer |
| %_forest | % of forests | class | % of broadleaf and coniferous tree cover within the buffer |
| %_open | % of open vegetated habitat | class | % of herbaceous vegetation, moors, heathland, sclerophyllous vegetation, marshes, and peat bogs within the buffer |
| frac | fractality | patch | complexity of the sampling patch |
| cai | core area index | patch | area of the sampling patch minus the pixel touching an other patch (percentage) |
| ed | edge density | class | sum of all meadow edges divided by the area of the raster (metres per hectare) |
| iji | interspersion and juxtaposition index | index | intermixing of meadow within other habitat classes (percent) |
| clumpy | clumpiness | class | calculate the disaggregation of the meadows |
| division | division | class | probability that two meadow points are not in the same patch |
| shdi | Shannon diversity index of habitat | landscape | descriptor increasing according to the number of landscape classes and evenness of these classes |

### 2.4 Protist community

#### 2.4.1 DNA extraction, PCR, sequencing

DNA was extracted from 0.25 g of soil samples in triplicates with the MoBio PowerSoil DNA extraction kit (Mo Bio Laboratories, Carlsbad, CA, USA) following the manufacturer’s instructions. The V4 region of the 18S rRNA gene was then amplified using the general eukaryotic primers `TAReuk454FWD1` and `TAReukREV3` (Stoeck et al., 2010). PCR amplification details can be found in Seppey et al. (2020). Paired-end reads (2 * 300 bp) were then sequenced from an Illumina MiSeq sequencer at the University of Geneva (Molecular Systematics & Environmental Genomics Laboratory) as in Seppey et al. (2020). Sequences are available on the European Nucleotide Archive under the project PRJEB30010 (ERP112373).

#### 2.4.2 Bioinformatic pipeline

Taken together, all samples provided 128’581’740 raw reads that were analysed with a pipeline based on `VSEARCH` v. 2.15.2 (Rognes et al., 2016). The reads were first merged with `Flash` v. 1.2.11 (Magoc and Salzberg, 2011) before removing the sequences shorter than 250 nucleotides. Reads with a quotient between expected error and sequence length below 0.01 as well as chimaeras were discarded. The 23’486’785 reads left (18%) were then clustered into 1’215’009 OTUs with the `Swarm` algorithm using the fastidious option (v. 3.1.0; Mahé et al., 2022).

The taxonomic assignment of the OTUs was performed in two steps. The first part was aiming to identify and remove the OTUs not belonging to protists (i.e. Prokaryotes, Metazoa, Embryophyta and Fungi). This was done by using the best alignment from a pairwise local alignment (`blastn` v. 2.9.0; Camacho et al., 2009) of the OTUs dominant sequences on the pooled PR^2^ 4.12.0 (Guillou et al., 2013) and Silva v. 138 (non-redundant at 99 %; Pruesse et al., 2007) databases. The second part consisted of the final assignment of the remaining OTUs by aligning their dominant sequences against the PR^2^ database v. 4.12.0 using the best alignment of a global pairwise alignment (`ggsearch36` v. 36.3.8; Pearson, 2000). Prior to the global pairwise alignment, the PR^2^ database was trimmed according to the primers used for the sequencing after a multiple sequence alignment performed with `ClustalW` v. 2.1 (Larkin et al., 2007).

From the 178 plots, 4 were sampled twice and 13 three times during the sampling period. For each of these plots, OTUs abundance was estimated as the median of the number of sequences retrieved from the different samples. In addition, 16 plots with a low number of sequences were removed by using the breaking point of a piecewise linear model on the sorted log of the number of sequences per sample (observed threshold = 6047 sequences) (Figure S3). In total, 162 samplesplots were kept for further analyses. We then discarded the OTUs with near-zero variance (ratio between the two most frequent OTU abundance below 95/5, and unique number of sequence within a OTU below 10% of the number of samples; function `nearZeroVariance`, package `caret` v. 6.0-86; Kuhn, 2020). The final dataset included 162 samplesplots and 3992 protist OTUs.

### 2.5 Statistical analyses

To assess local α diversity, the Shannon diversity index was calculated for each community. To test the importance of landscape structure for soil protist α diversity, a random forest algorithm was used to model the response of soil protist Shannon diversity to landscape and environmental descriptors. Prior to the modelling, all landscape and environmental descriptors were scaled to a standard deviation of 1 and centred to a mean of 0. Several runs of random forest were conducted within a 10-fold cross-validation stratified according to the response variable, repeated three times. The models used 500 trees and a maximum depth of 125 nodes (default parameters of the function `train`, package `caret` v. 6.0-86; Kuhn, 2020). The number of variables per split (‘mtry’ = 2, 14, 27; hyperparameter tuning) was selected to optimise model performance (based on the best R^2^). The importance of each descriptor was estimated as the mean decrease in accuracy after permuting the target descriptor (function `varImp`, package `caret` v. 6.0-86; Kuhn, 2020). To allow comparisons among models, the importance of each descriptor was weighted by the R^2^ of the model. To assess the directions of the effect, partial dependence plots (Goldstein2015, function `partial`, package `pdp` v. 0.7.0 Greenwell, 2017) were computed for each descriptor. Partial dependence plots summarise the marginal effect that an environmental or landscape descriptor has on the predicted values of each diversity metric (i.e. effect of a target descriptor when all others are kept constant). This procedure was repeated for each neighbourhood window.

Changes in protist community composition among sites (β diversity) were measured using abundance-based Bray-Curtis dissimilarity following the approach of Baselga et al. (2013). We assessed the rate of distance-decay of the protist communities as the slope of a linear regression on the relationship between protist Bray-Curtis dissimilarity and geographic distance, both log-transformed. Because of the non-independence of pairwise comparisons, we used a matrix permutation test to assess the statistical significance of the distance-decay slope. Specifically, the rows and columns of the protist Bray-Curtis dissimilarity matrix were permuted 1,000 times, and the observed slope was compared with the distribution of values in the permuted datasets. We also tested whether the environmental and landscape descriptors (for each of the five neighbourhood windows) were responding to geographic distance using the same permutation procedure.

To test whether landscape structure drives changes in protist community composition, we first performed a RDA on the protist community dissimilarity matrix using landscape, environmental and spatial descriptors. The spatial descriptors consisted of a selection of 16 of 80 principal coordinates of neighbourhood matrix (PCNM) calculated from the euclidean distances between plots (Borcard and Legendre, 2002). The PCNM descriptors showing the strongest correlations to the protist Bray-Curtis dissimilarities were selected by stepwise selection (function `ordistep`, package `vegan` v2.6-2; Oksanen et al., 2022). We used a `PERMANOVA` (function `anova.cca`, package `vegan` v2.6-2; Oksanen et al., 2022) to test the significance of marginal effect of each variable included in the RDA. Secondly, we tested the variation explained by each of the three sets of variables through variation partitioning (Borcard et al., 1992, function `varpart`, package `vegan` v.2.6-2; Oksanen et al., 2022).

The bioinformatic pipeline and all statistical analyses can be found on https://gitlab.com/cseppey/milan. All statistical analyses were performed in `R` 4.2.2.

## 3 Results

### 3.1 Protist community composition

Soil protist diversity was dominated by Alveolata and Rhizaria, which represented 37% and 33% of the sequences, respectively. Figure S4 provides further information about protist community composition.

### 3.2 Influence of landscape structure on soil protist diversity

Landscape structure had a significant effect on protist Shannon diversity. The amount of variation in protist Shannon diversity explained by landscape descriptors was greater than the one explained by environmental descriptors when using larger neighbourhood windows (1000 m to 2000 m around plot centre) and lower when using smaller neighbourhood windows (100 m and 500 m: Figure 2A). The percentage of open vegetated habitats around a plot explained the highest amount of variation in protist Shannon diversity among all landscape descriptors (Figure 2B), whereas patch level descriptors (patch area, patch core area, and patch fractality) explained the lowest amount. The three landscape descriptors that explained the largest part of soil protist Shannon diversity were the percentage of open vegetation in 500 m neighbourhood windows, the percentage of meadows (1000 m) and edge density (1000 m). Specifically, protist α diversity was positively correlated with the percentage of open vegetation (500 m) and followed a unimodal relation with the percentage of meadows (1000 m) and edge density (1000 m) (Figure 2C). The importance and direction of the effect of each landscape descriptor on Shannon diversity can be found in Figure S5.

**Figure 2:**
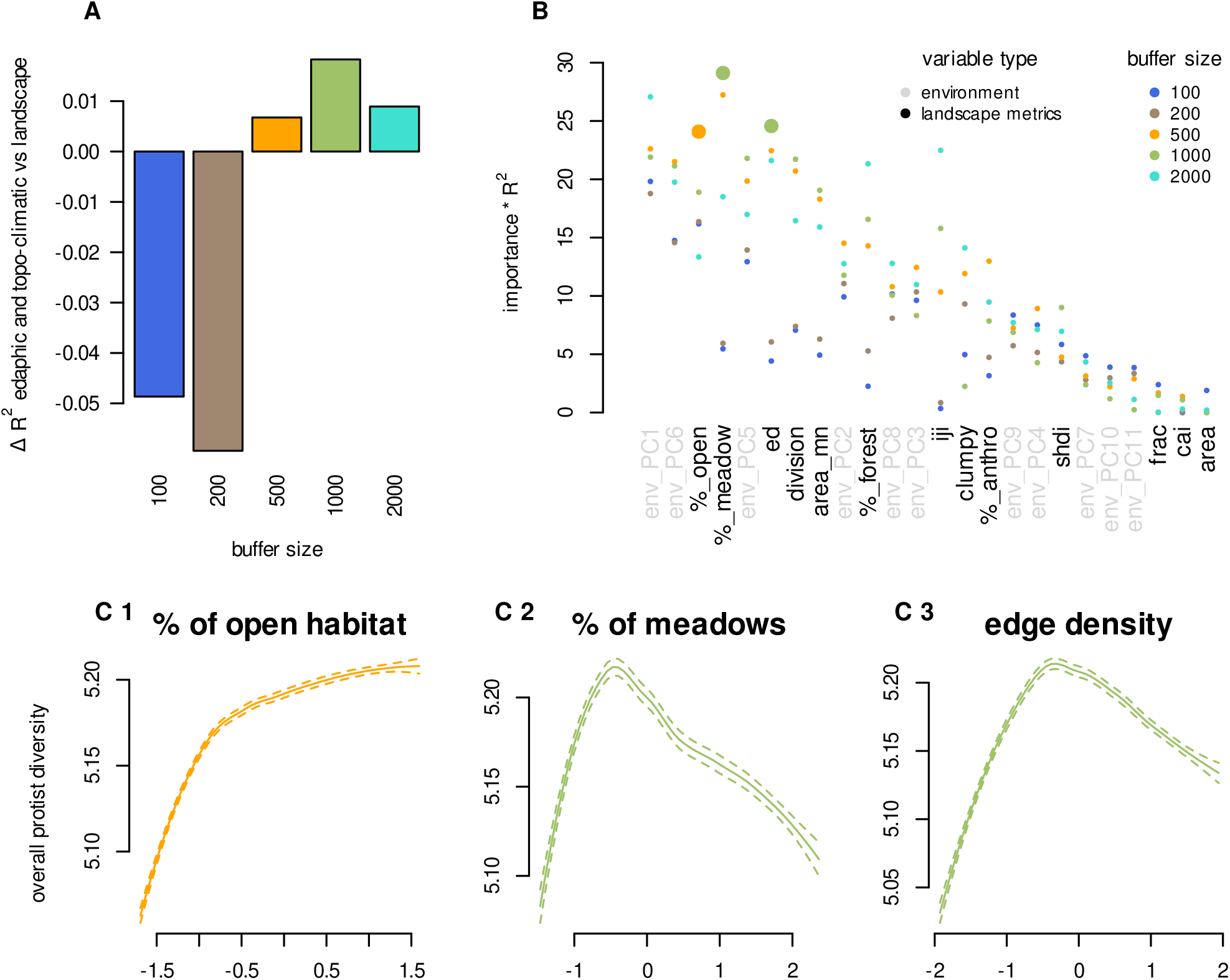
Effect of landscape structure on soil protist Shannon diversity. (A) Difference between the variation in total protist diversity explained by landscape descriptors versus environmental descriptors in random forest models (R^2^_landscape_ - R^2^_environment_), (B) Variability in the importance of landscape and environmental descriptors for the description of overall soil protist Shannon diversity in random forest models. The importance has been weighted by the R^2^ of the model to allow comparisons among models. Landscape descriptors are shown in black, and environmental descriptors in grey. The three landscape descriptors with the highest importance are highlighted (percentage of open vegetated habitats, edge density, and percentage of meadows). (C) Partial dependence plots showing the trends of soil protist Shannon diversity along the three most important landscape descriptors. For each sub-figure, the y-axis reflects the effect that the target landscape descriptor has on the predicted values of the response variable while keeping all other descriptors constant. The x-axis represents the centred and scaled values of the three landscape descriptors that best explain protist Shannon diversity. Colours indicate the most important (*importance* * *R*^2^) neighbourhood window sizes.

### 3.3 Changes in protist community composition

We found a significant positive relationship between protist community dissimilarity and geographic distance among plots (slope = 0.015, P < 0.01: Figure 3A). In addition, we found significant positive relationships between the dissimilarity of environmental or landscape descriptors and geographic distance among plots (Figure 3B). It is noteworthy that the slopes of environmental descriptors (slope = 0.106) were less steep than the ones calculated for landscape descriptors. The slopes of the landscape-based DDR were also steeper for larger neighbourhood window sizes (100 m: 0.111; 200 m: 0.133; 500 m: 0.180; 1000 m: 0.194; 2000 m: 0.269)

**Figure 3:**
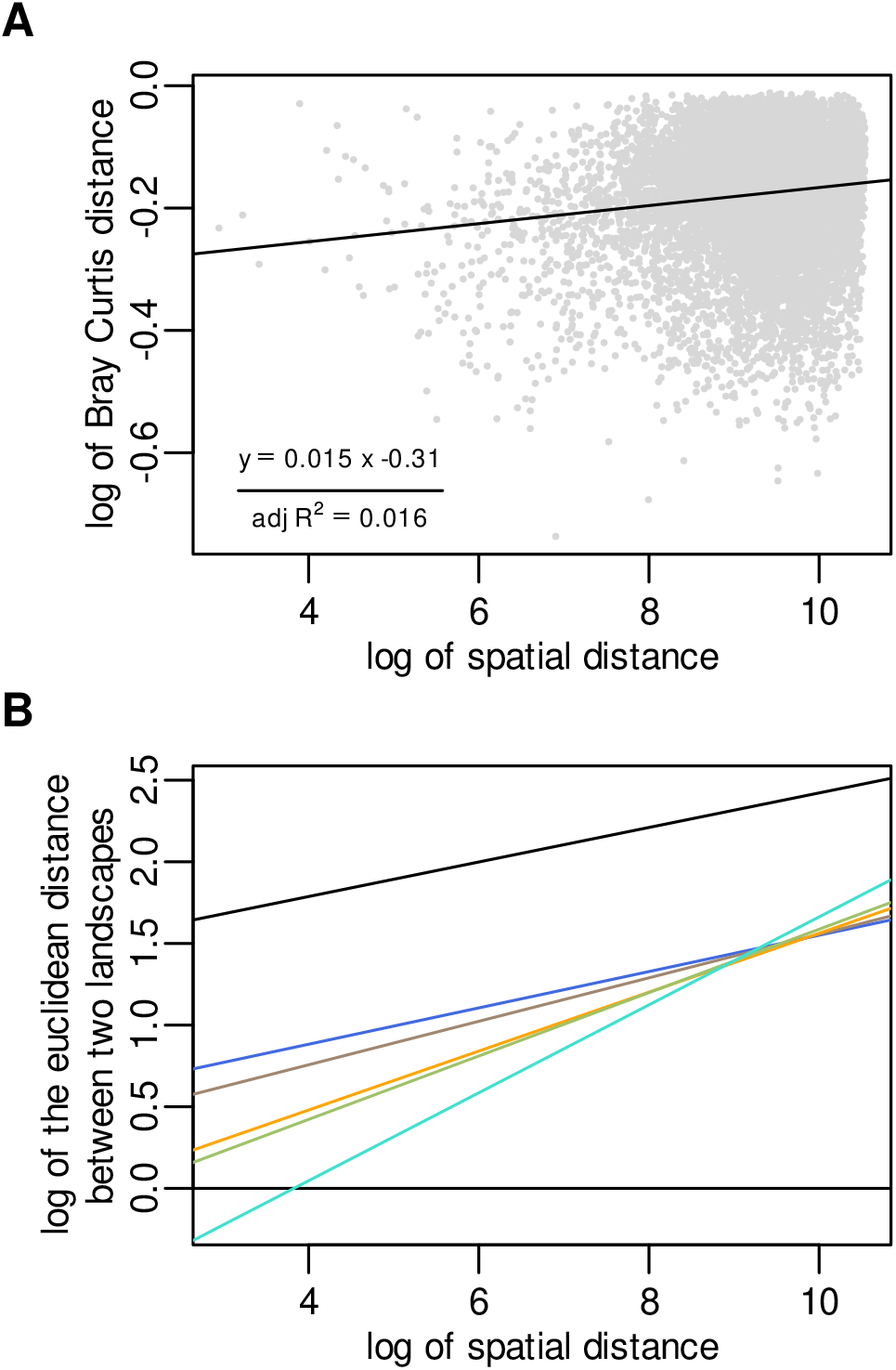
Distance-decay curves for the (A) protist communities and (B) landscape and environmental descriptors. All Distance-decay curves are shown in log-log scale for optimal visualisation. (B) The different lines correspond to distance-decay relationships for environmental (black) and landscape descriptors calculated in neighbourhood windows of 100 m (blue), 200 m (brown), 500 m (orange), 1000 m (green) and 2000 m (cyan). All slopes were significantly different from a null model obtained by permuting 1000 times the samples of the dependent variable with p-values ≤ 0.001.

Redundancy analyses showed that landscape, environmental and spatial descriptors jointly influenced protist β diversity (Figure 4). The amount of variation in protist β diversity explained by landscape descriptors was comparable to the one explained by environmental or spatial descriptors. It is also noticeable that the landscape effect increased with the size of the neighbourhood window, even explaining a slightly higher portion of the variation in protist community composition than spatial or environmental variables when calculated in larger neighbourhood windows (2000 m, Figure 4). In addition, `PERMANOVA` analyses showed that the landscape variables that significantly contribute to explaining changes in protist β diversity varied among scales (Table 2). For instance, the percentage of anthropic, forested or vegetated open area played a major role at the smaller scales (100 m and 200 m windows) while landscape descriptors such as clumpiness or the Shannon diversity index of habitat were only significant when calculated in larger windows (500 m to 2000 m windows).

**Figure 4:**
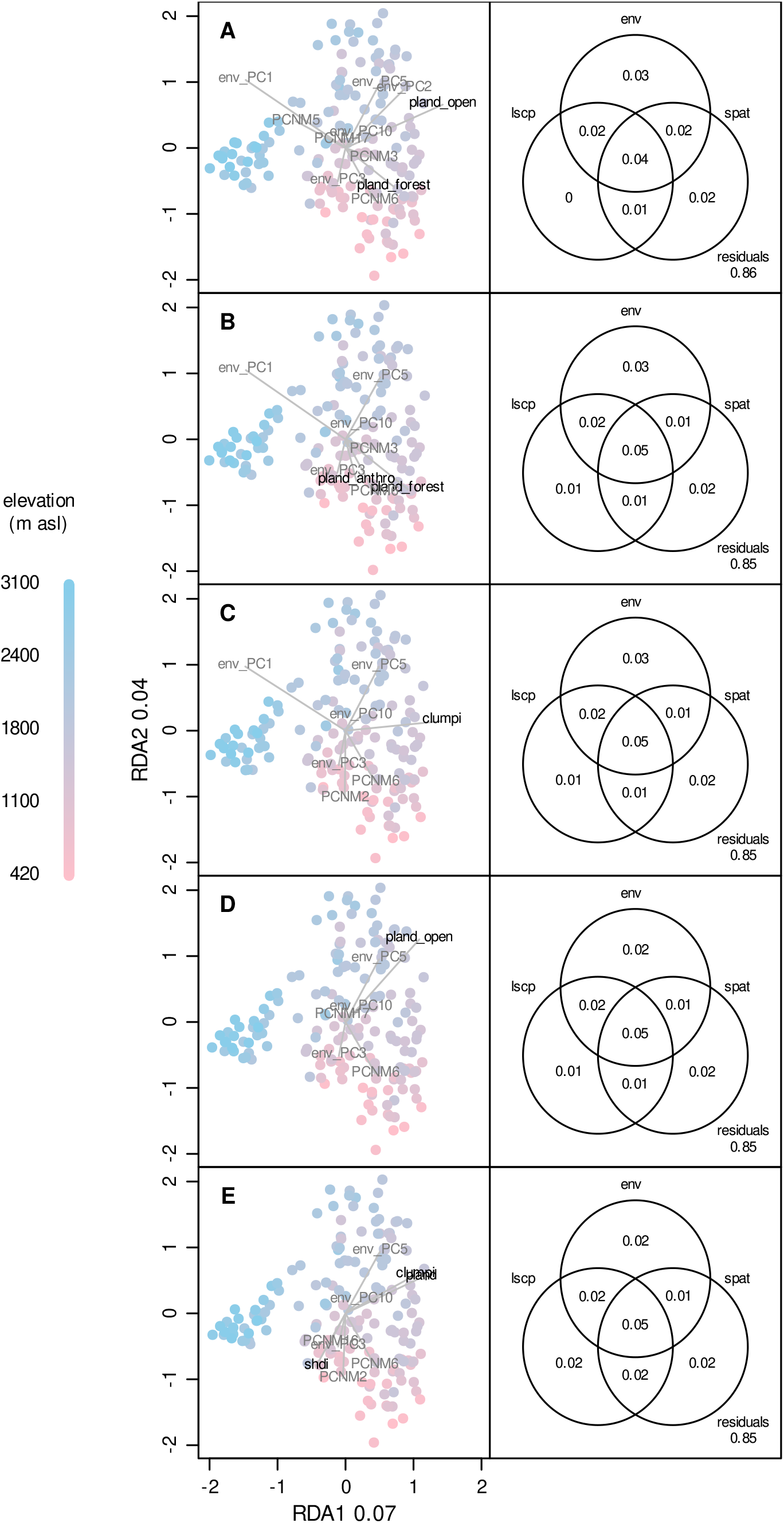
Redundancy analyses showing the effect of landscape (lscp), environmental (env), and spatial variables (PCNM, spat) on protist communities. The panels on the left show the full RDAs with arrows representing the contribution of significant variables (see Table 2). The five RDAs were done using landscape variables calculated in different neighbourhood sizes (A: 100 m; B: 200 m; C: 500 m; D: 1000 m; E: 2000 m). The variance on the two first axes range between 0.066 and 0.068 and 0.037 and 0.040, respectively. The total variance explained by the RDAs range between 0.14 and 0.15 (adjusted-R2). The panels on the right show the partitioning of the variation in protist communities among landscape, environmental and spatial variables. The significance of the different fractions are given in Table S3.

**Table 2:**
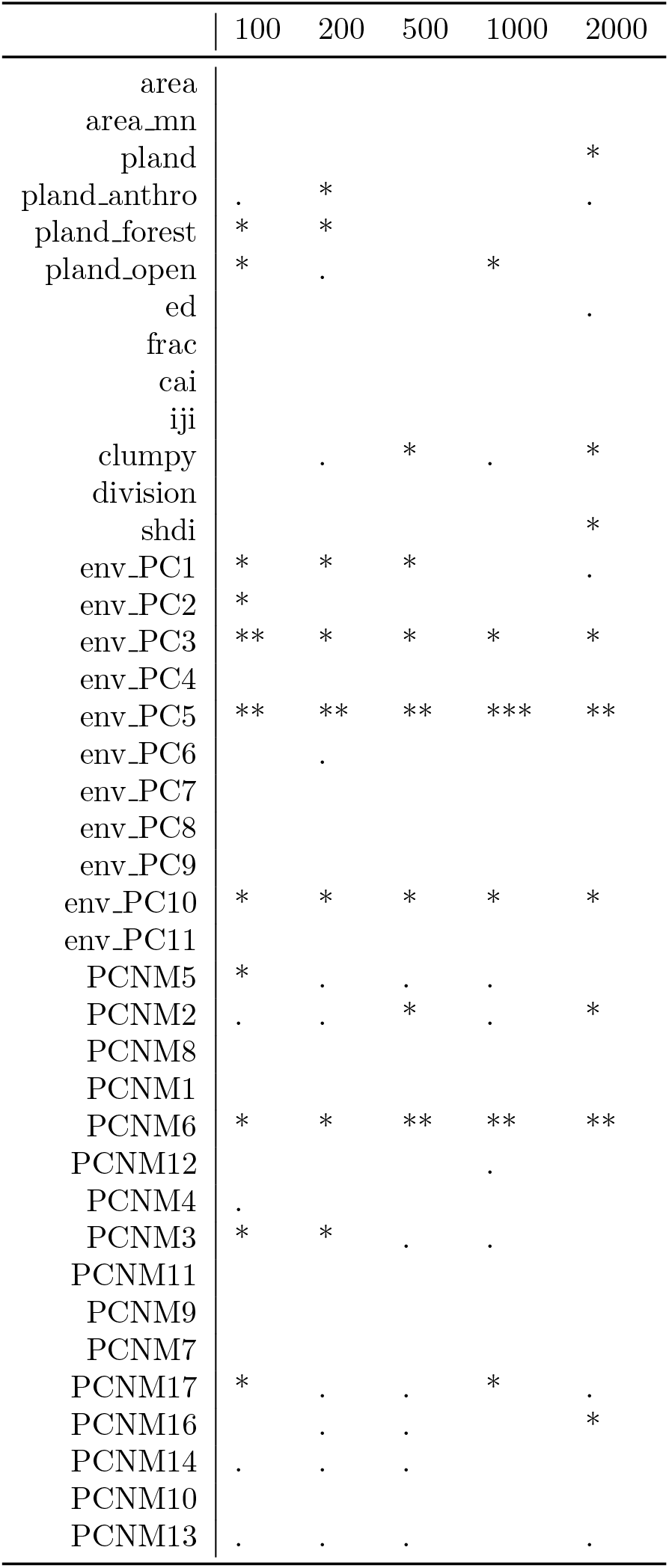
Summary statistics of permutation-based analysis of variance (PERMANOVA). The significance of the marginal effect of each variable included in the RDA is shown. Stars indicate significant p-values: “***” ≤ 0.001, “**” ≤ 0.01, “*” ≤ 0.05, “.” ≤ 0.1.

## 4 Discussion

We showed the importance of landscape structure in explaining protist α and β diversity. This adds to the body of evidence showing the importance of spatial environmental factors such as topography, and climate (Seppey et al., 2020) in structuring protist communities and provides new insights on the potential effects of land-use change. Landscape descriptors show comparable importance to environmental and spatial ones, which establish them as a promising tool for a wide range of spatial analyses. As expected, the effect of landscape structure on both protist α and β diversity was scale-dependent and maximal for neighbourhood windows between 500 m and 2000 m. These results confirm that local protist community composition strongly depends on landscape-scale assembly processes such as dispersal from the regional pool of taxa.

### 4.1 Landscape structure influences local protist α diversity

Our results highlight the key role of landscape structure for soil protist α diversity in alpine meadows. Landscape composition descriptors (e.g. percentage of open vegetated habitats or meadows around each plot) had the most important effect on soil protist diversity. On the contrary, landscape configuration descriptors (e.g. division, interspersion and juxtaposition index), although important, contributed less. Interestingly, patch-scale descriptors had a consistently low influence on soil protist diversity and, therefore, cannot explain the observed changes in protist diversity. Overall, these results agree with the ‘habitat amount hypothesis’ that suggests that the proportion of habitat type in the landscape is more important than its spatial distribution (Fahrig et al., 2015; Laine and Hanski, 2006). These conclusions are similar to those of previous works on the distribution of fungi in old forests (Mennicken et al., 2020), suggesting that similar mechanisms impact protists and fungi following land-use change. While these convergent lines of evidence suggest that these conclusions are valid for all soil microbes, they are in strong contradiction with studies focusing on plants and animals, where patch size and shape have strong effects on local population dynamics (Prevedello and Vieira, 2010; Reed, 2004). This difference between macro- and microorganisms can be explained by differences in population size. Indeed, for the same area, the population size of microbes is larger by several orders of magnitude than that of macroorganisms. As such, population size fluctuations and genetic drift (and other population processes) are less likely to be problematic for microbes at the scale of a meadow.

In our study, a high proportion of open vegetated habitats around the plot had a positive effect on local diversity whereas a high proportion of artificial habitats (built-up and cultivated) had a negative one (Figure S5). Maximal protist diversity was observed for intermediate proportions of forest in the surrounding landscape (Figure S5). This can be explained by three different processes. First, protist communities in forests have been shown to be less diverse than those in meadows (Seppey et al., 2017). Then, open vegetated habitats are not homogeneous and include a large variety of ecosystem types (herbaceous vegetation, moors, heathland, sclerophyllous vegetation, marshes, and peat bogs), each one having its specific protist communities. For this reason, open vegetated habitats as a whole are more likely to host a higher diversity than forests (more homogeneous) or artificial habitats. Assuming similar niche-based filtering or dispersal among habitats, these differences in protist diversity among habitats are sufficient to explain the observed patterns. However, another potential explanation is that the closed structure of forest and artificial habitats hampers protist dispersal, which is likely caused by wind (Wilkinson et al., 2012). If true, fewer taxa should be able to colonise a meadow plot from the forest or artificial habitats than from open vegetated habitats. It follows that differences in dispersal alone could explain the observed patterns. Finally, the different environmental conditions such as soil type and moisture can hamper species from the forest or artificial habitat to colonise the meadow which would suggest a stronger filtering through niche processes of species adapted to forest or artificial habitats. These three potential explanations are not mutually exclusive and interactive effects are likely.

Despite the higher influence of landscape composition on protist α diversity, our results also point towards strong edge effects (Figure 2C). Edges are hotspots of biodiversity because of the presence of species associated with different habitats alongside ecotonal species (Descombes et al., 2017). They are also an important source of dispersal to neighbouring habitats (Molnar et al., 2001). In microbial ecology, only a few studies have investigated community composition in transition zones between habitats. For example, Boeraeve et al. (2019) showed that both distance to the forest edge and edge orientation to the south influence mycorrhizal communities of trees (*Alnus glutinosa*). In our study, soil protist diversity was maximal at intermediate edge density. A low edge density means a lower likelihood that colonists from adjacent habitats settle down in the meadow. Ecotonal specialists might also be lacking when edge density is too low. The lower local protist diversity at high edge density can tentatively be explained by two mechanisms. On the one hand, high edge density might favour a few generalist species over specialist taxa. Generalists can survive in various habitats and are more likely to benefit from highly mixed landscape habitat mosaics. On the other hand, edge density was positively correlated with the area of urban habitat (Spearman correlation P < 0.001, ρ= 0.44). As these habitats have a negative contribution to local diversity, soil protist diversity is lower than when only habitats with positive contributions to diversity are present. In this case, edge density would reflect a compositional effect rather than a true edge effect.

Altogether, protist α diversity in the Alps is favoured by a diversified landscape, containing equilibrated amounts of meadows and forests, but also to some extent artificial habitats. This stands in contrast with current tendencies towards increasing urbanisation and meadow recolonisation by forests as a result of agricultural abandonment (Spiegelberger et al., 2006). Diversified protist communities are key in many soil ecosystem services, such as carbon storage and plant production (Geisen et al., 2019). The disappearance of traditional lifestyles in the Alps (Spiegelberger et al., 2006) can be expected to modify the structure and functioning of the soil microbial foodweb. Such modifications can lead to more negative impacts on Alpine meadows than previously assessed based on macroorganisms alone (Spiegelberger et al., 2006). Overall, maintaining sufficient landscape diversity in the Alps while avoiding excessive fragmentation is thus likely to have a positive effect on meadow soil protist diversity and functionality as observed in macroorganism groups such as Orthoptera (Essl and Dirnboeck, 2012), or plants and pollinators in general (Jones et al., 2019).

### 4.2 Landscape structure modulates the rate of distance-decay in protist β diversity

We found a significant distance-decay relationship in protist community (dis-)similarity (Figure 3A) as already observed in different environmental contexts (Lentendu et al., 2018; Macingo et al., 2019). This distance-decay can in part be explained by changes in environmental conditions (niche-based processes), especially soil moisture (env_PC5) and organic carbon content (env_PC3) which were the most significant environmental descriptors in the RDAs (Figure 4) and by dispersal effect due to a decrease in propagule flow among habitats with distance. Importantly, our results highlight that landscape structure has a key effect on the rate of distance-decay. Landscape structure indeed explained a similar portion of the variation in protist community composition as environmental and spatial variables (Figure 4). Landscape distance-decay was also significant, showing that two plots far apart tend to have more different landscape structure than two plots close to each other (Figure 3B). Furthermore, the effect of landscape structure on the rate of distance decay was dependent on the size of the neighbourhood window used to calculate landscape descriptors. Indeed, the slopes of the DDRs (Figure 3B) and the importance of landscape effects increased with the size of the neighbourhood window (Figure 4). In addition, different landscape descriptors correlates with protist dissimilarity depending on the size of neighbourhood window considered (Table 2) suggesting scale-dependent effects of landscape structure on the rate of distance-decay.

Overall, our results suggest that landscape structure modulates the rate of distance-decay by influencing niche-based and dispersal processes. Three main mechanisms can be evoked here: dispersal excess (mass effect, Leibold et al., 2004), environmental filtering (species sorting), and dispersal limitations. Dispersal excess or mass effect can be expected over short distances (where dispersal limitations are unlikely) and among habitats having different environmental characteristics (source and sink habitats) (Mouquet and Loreau, 2003; Leibold et al., 2004; Urban, 2006; Logue et al., 2011). This fits with the significant effect of percentage of forest and anthropic habitats in neighbourhood windows of 100 m or 200 m radii on protist community dissimilarity. Under mass effects, two meadows surrounded by similar forest habitats will host an important number of forest species even if those species are maladapted to local conditions. This will result in a lower β diversity than expected based on local environmental differences or geographic distance alone. Species sorting results in a matching between the environmental conditions and community composition and is expected in the absence of dispersal limitation or excess. Landscape structure determines the taxonomic composition and size of the landscape pool of taxa that can potentially colonise a focal plot. Two plots with similar environmental conditions will impose a similar filtering on the landscape pool of potential colonists. However, if the composition of those pools differ (e.g. because of different landscape structure), environmental filtering will select or filter different species thereby impacting β diversity. In agreement with this idea, the observed effect of the percentage of open habitats (1000 m) and meadows (2000 m) can be related to changes in the composition of the landscape pool of potential colonists. Finally, landscape structure has been shown to influence the rate of organism dispersal among habitat patches in several taxonomic groups such as plants (Auffret et al., 2017), mammals (Hämäläinen et al., 2019), and arthropods (Bonte et al., 2006). In our case, low clumpiness of meadows can increase inter-patch distances thereby increasing dispersal risks and reducing movements among patches (Fahrig, 2007). Furthermore, landscape composition can facilitate or hamper dispersal rates among habitat patches. For instance, large patches of low vegetation such as in moors and heatlands or sclerophylous vegetation can facilitate dispersal by wind whereas the closed structure of forest patches might hamper dispersal by wind. However, how landscape structure influences the dispersal of protists likely depends on the dispersal mode of specific taxa, where important differences between free living taxa (expected to disperse by wind) and parasites (depending on host movement) can be expected. Generally, the precise mechanisms responsible for the patterns observed in this study need to be confirmed by experimental studies.

Overall, our results highlight the importance of dispersal from surrounding habitats for protist community composition. Microbial community dynamics should thus be considered at the landscape-scale using a metacommunity perspective (Beisner et al., 2006; Cadotte, 2006; Leibold et al., 2004) which has important consequences for land-use management in the Alps and in general. Indeed, changes in landscape structure are likely to modify assembly processes, thereby impacting the composition, functionality and resilience of soil protist communities (Saleem et al., 2019). As soil protists are key players in soil food webs (Xiong et al., 2018), these changes can potentially have a cascading impact on ecosystem functioning and services such as fertility as well as carbon sequestration and storage (Jassey et al., 2022). It is thus likely that the ongoing agricultural abandonment and urbanisation in the Alps (Vannier et al., 2016) will lead to local and regional decreases in protist diversity in meadow soils with potentially important consequences for the functioning of these ecosystems.

### 4.3 Future developments of microbial landscape ecology

While the present study and several other recent works (Mennicken et al., 2020; Mony et al., 2021, 2022) provide a strong basis to further develop the field of microbial landscape ecology, much remains to be done. One of the most important questions arising from our work is the actual rate of dispersal (excess or limitations) of protists and microbes in general from surrounding habitats (Frey, 2015). The influence of barriers to dispersal across different microbial taxa (Paz et al., 2021; Wilkinson and Mitchell, 2010) needs to be investigated in the future. Likewise, questions arise about the generalisability of our results. Would we observe a similar relationship linking landscape and protist diversity in other habitats or regions (e.g. tropical areas)? Would other microbial groups show similar patterns (e.g. bacteria, fungi, archaea)? This last question is worth investigating, as, for instance, Mennicken et al. (2020) showed that bacteria and fungi responded differently to landscape composition. Finally, while our results highlight the importance of landscape structure for local protist community diversity and composition, the functional consequences of those changes and their magnitude remain unclear. Similarly, the extent to which these changes influence other taxa, including other microorganisms and macroorganisms, needs to be investigated.

## Conclusion

Our results highlight the importance of landscape structure, especially changes in the composition of neighbouring habitats, in structuring soil microbial diversity in alpine meadows. Landscape structure in the larger neighbourhood windows (radii of 500 m to 2000 m) was as important as the commonly used edaphic parameters in describing protist α and β diversity. We showed that, like macroorganisms, soil protists are strongly impacted by changes in landscape structure, which is likely to have important consequences on ecosystem structure and functioning. Using landscape ecology approaches in microbial ecology studies can improve our understanding of the drivers of and capacity to predict spatial patterns of microbial diversity. Microbial landscape ecology is a promising avenue to improve our understanding of the impact of land-use changes on biodiversity and ecosystem functions.

## Acknowledgements

BF and CVWS acknowledge the WISNA program from the German Federal Ministry of Education and Research. BF acknowledges the project “FunShift” (project FO 1420/1-1) awarded by the German Research Foundation, DFG. EL acknowledges the program “Atracción de Talento Investigador” from the Community of Madrid (project 2017-T1/AMB5210) and the project Myxotropic VI PGC2018-094660-BI00 awarded by the Spanish Government.

## Authorship

The study was conceived by B.F. and C.V.W.S and designed with input from all authors. Sampling design, fieldwork, and laboratory analyses were carried out by A.G., E.Y., and D.S. C.V.W.S. carried out the bioinformatic and statistical analyses. B.F. and C.V.W.S. led the writing of the manuscript, all authors contributed to drafts of the manuscript and gave final approval for publication.

## Data Accessibility Statement

All DNA sequences are available on the European Nucleotide Archive project number: PR-JEB30010 (ERP112373). Edaphic variables are available from Dryad repository: https://doi.org/10.5061/dryad.fttdz08p7. Details about the topo-climatic data can be found at https://www.unil.ch/files/live/sites/ecospat/files/shared/PDF\_site/chclim25.pdf.

## Supplementary material

**Table S1:**
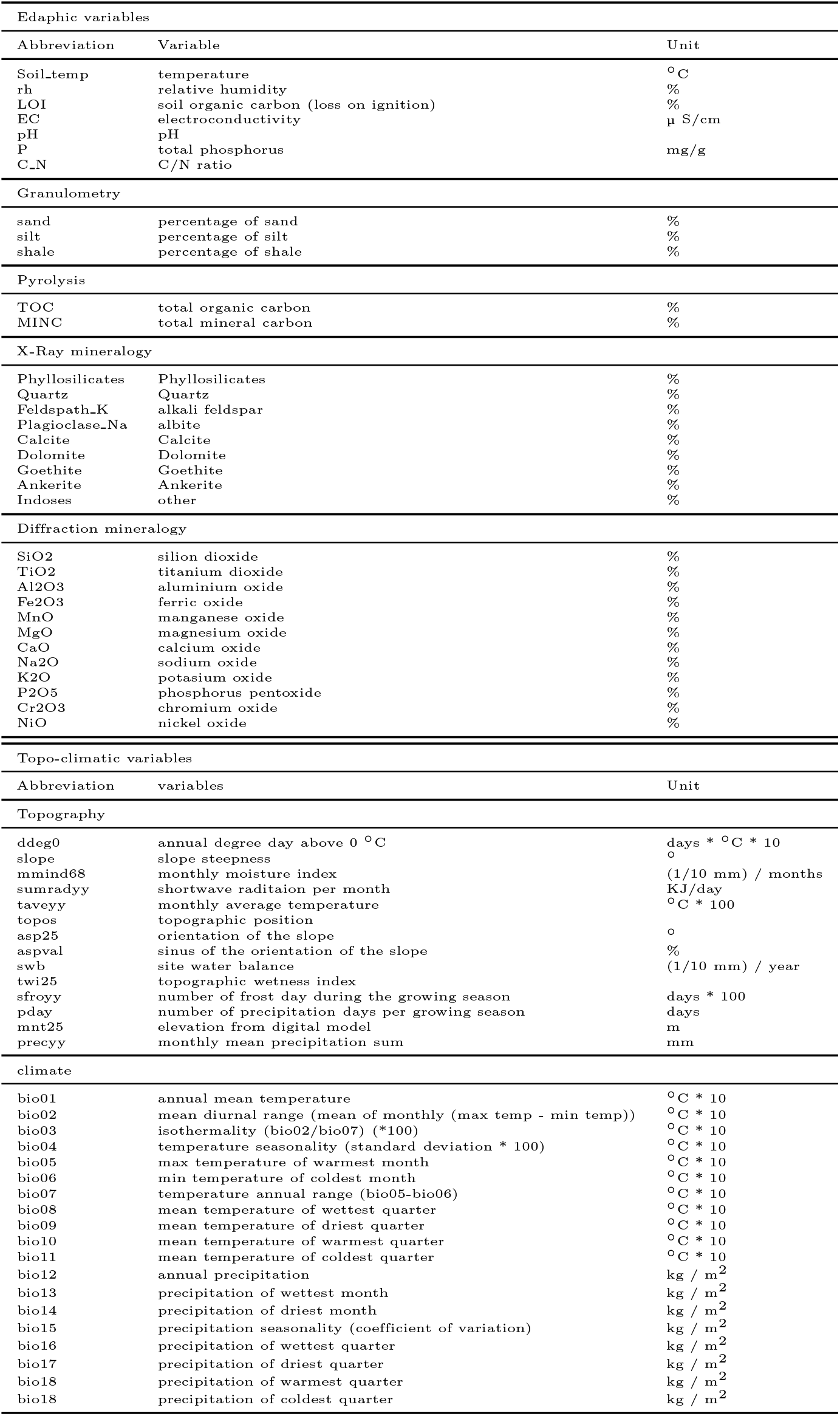
List of environmental descriptors and their abbreviations.

**Table S2:**
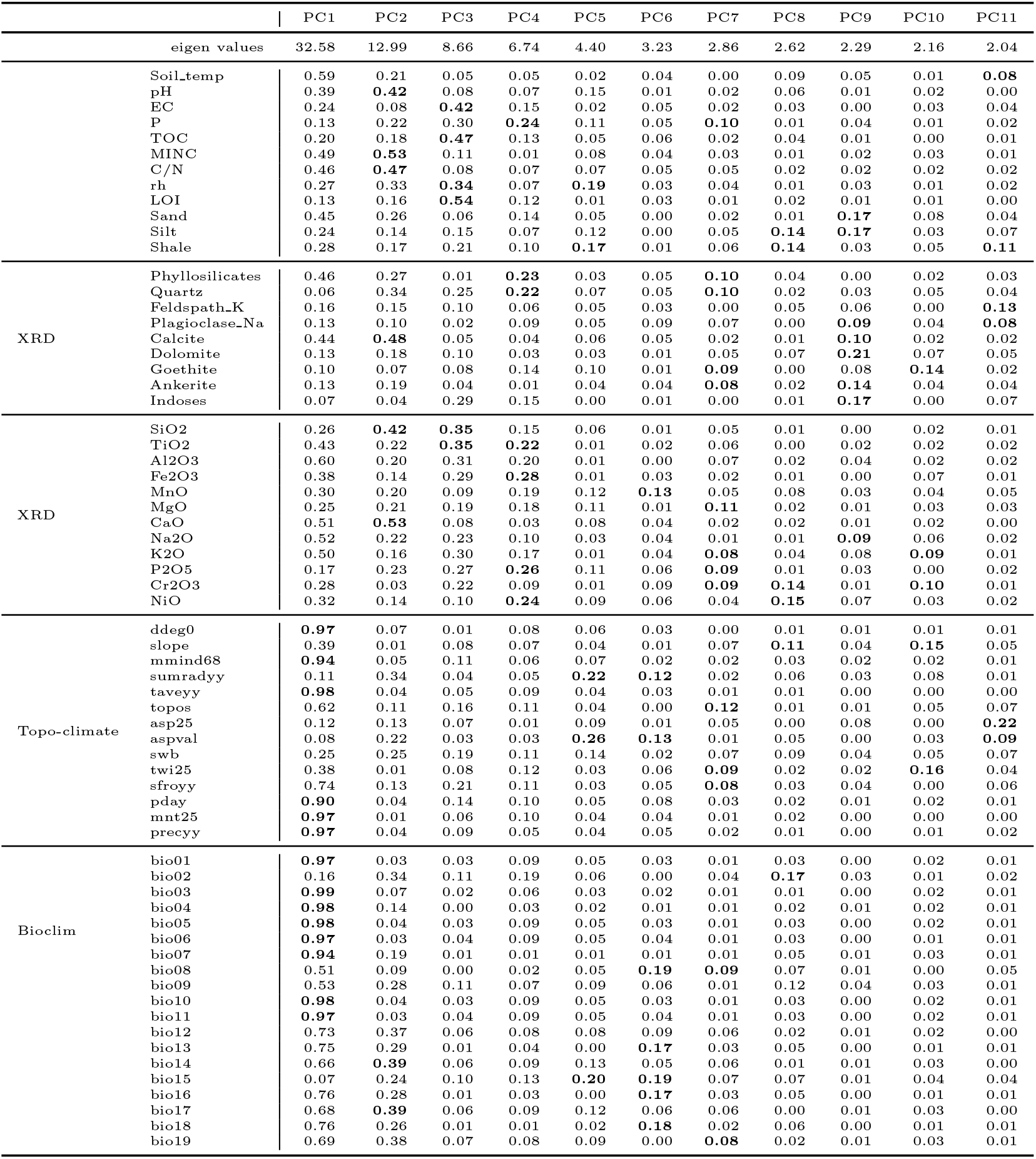
Contribution of the environmental descriptors measured from 162 meadow soils in the Swiss western Alps to the ten first axes of a principal component analysis. The contributions for each axis were calculated as the relative contribution of the loadings values multiplied by 100. The five highest values per axis are shown in bold.

**Table S3:**
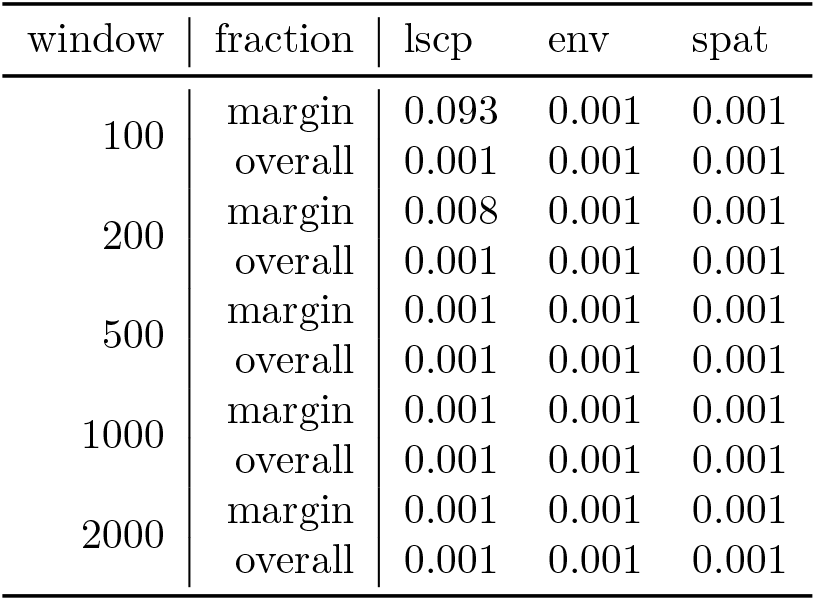
Significance of variation partitionings’ marginal and overall fraction between the Bray-Curtis dissimilarity of soil protist and landscape (lscp), environmental (env) and spatial (spat) descriptors.

**Figure S1:**
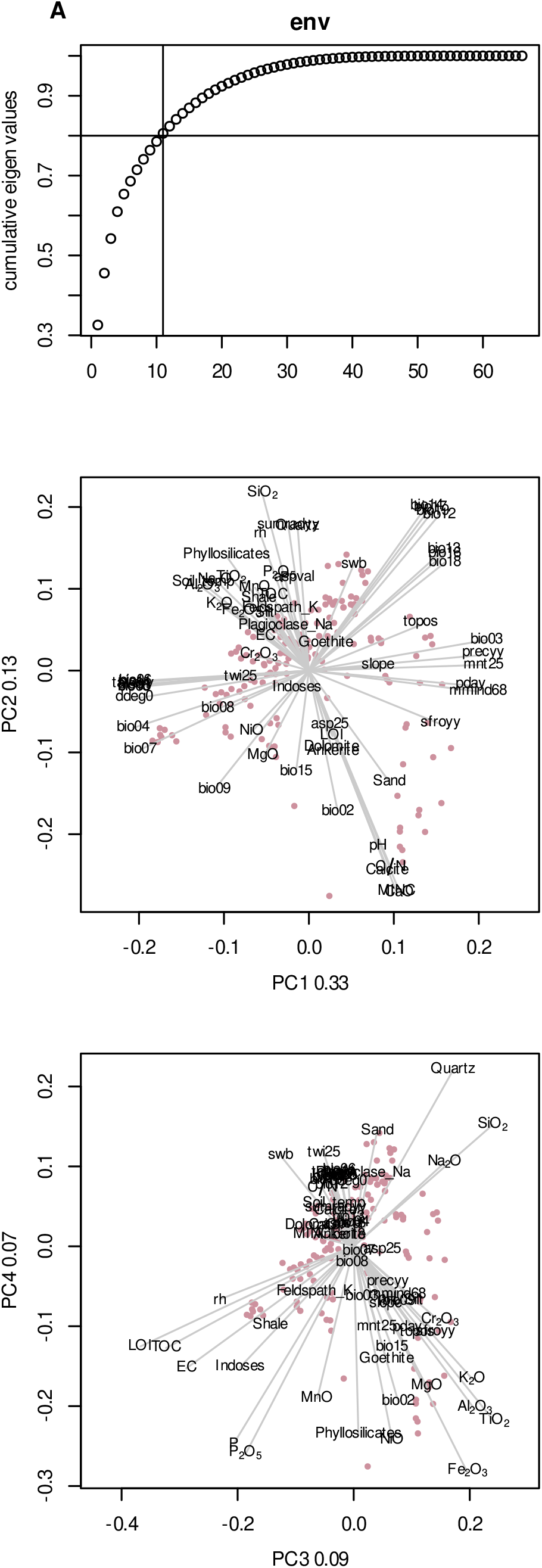
Principal component analysis (PCA) of the environmental descriptors measured from 162 meadow soil samples in the Swiss western Alps. Sub-figure A shows cumulative eigenvalues while sub-figures B and C show the four first PCA axes.

**Figure S2:**
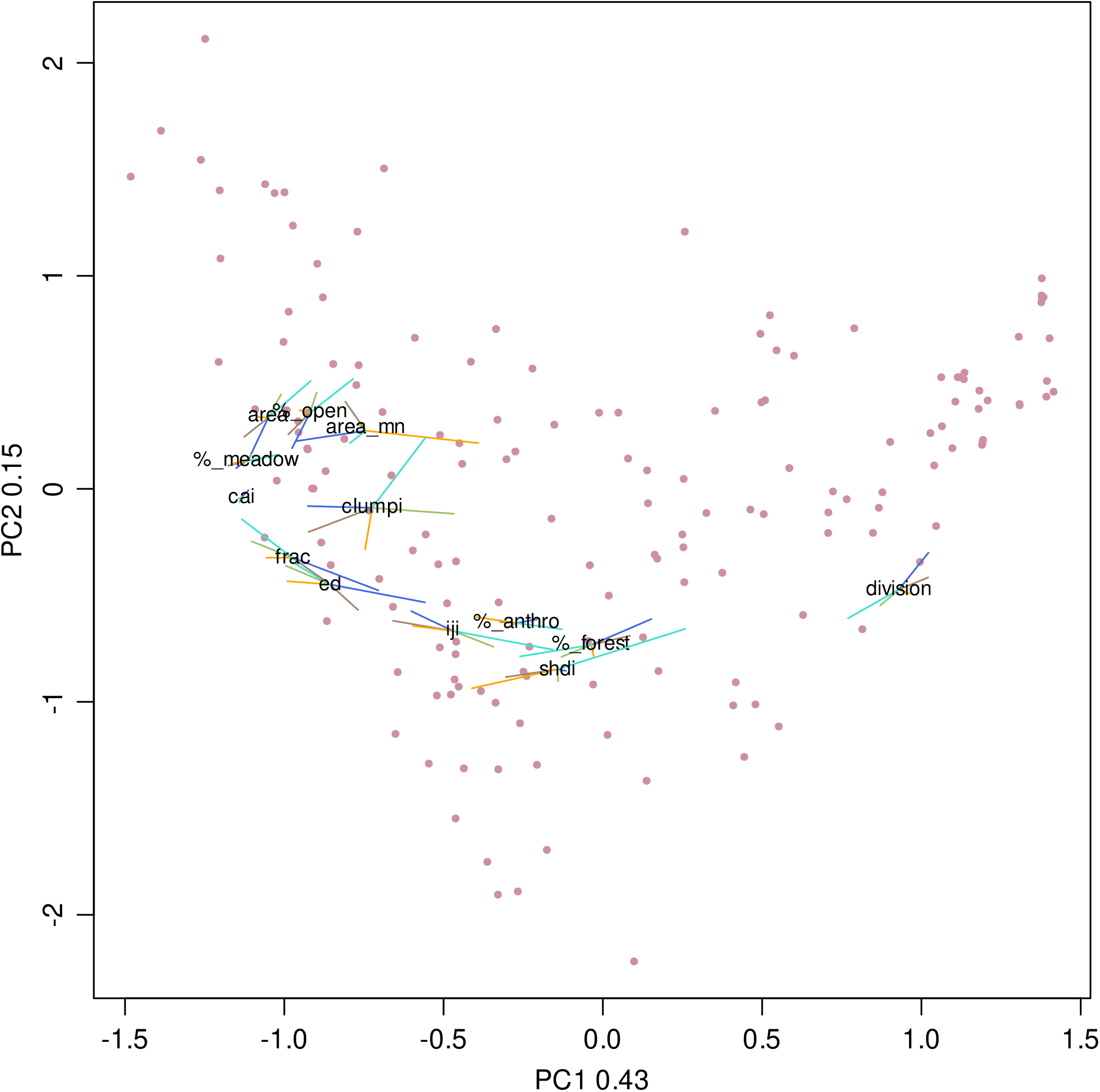
Principal component analysis (PCA) of 13 landscape descriptors measured around 162 meadows in the Swiss western Alps. Each landscape descriptor has been calculated in different neighbourhood windows: radius of 100 (blue), 200 (brown), 500 (orange), 1000 (green) and 2000 (cyan) metres around the sampling plots. Each landscape descriptor is represented by five points, one for each neighbourhood window (i.e. tip of the colour lines). Descriptor’s names are placed at the centre of gravity of these five points and represent the average effect of a descriptor across all neighbourhood windows.

**Figure S3:**
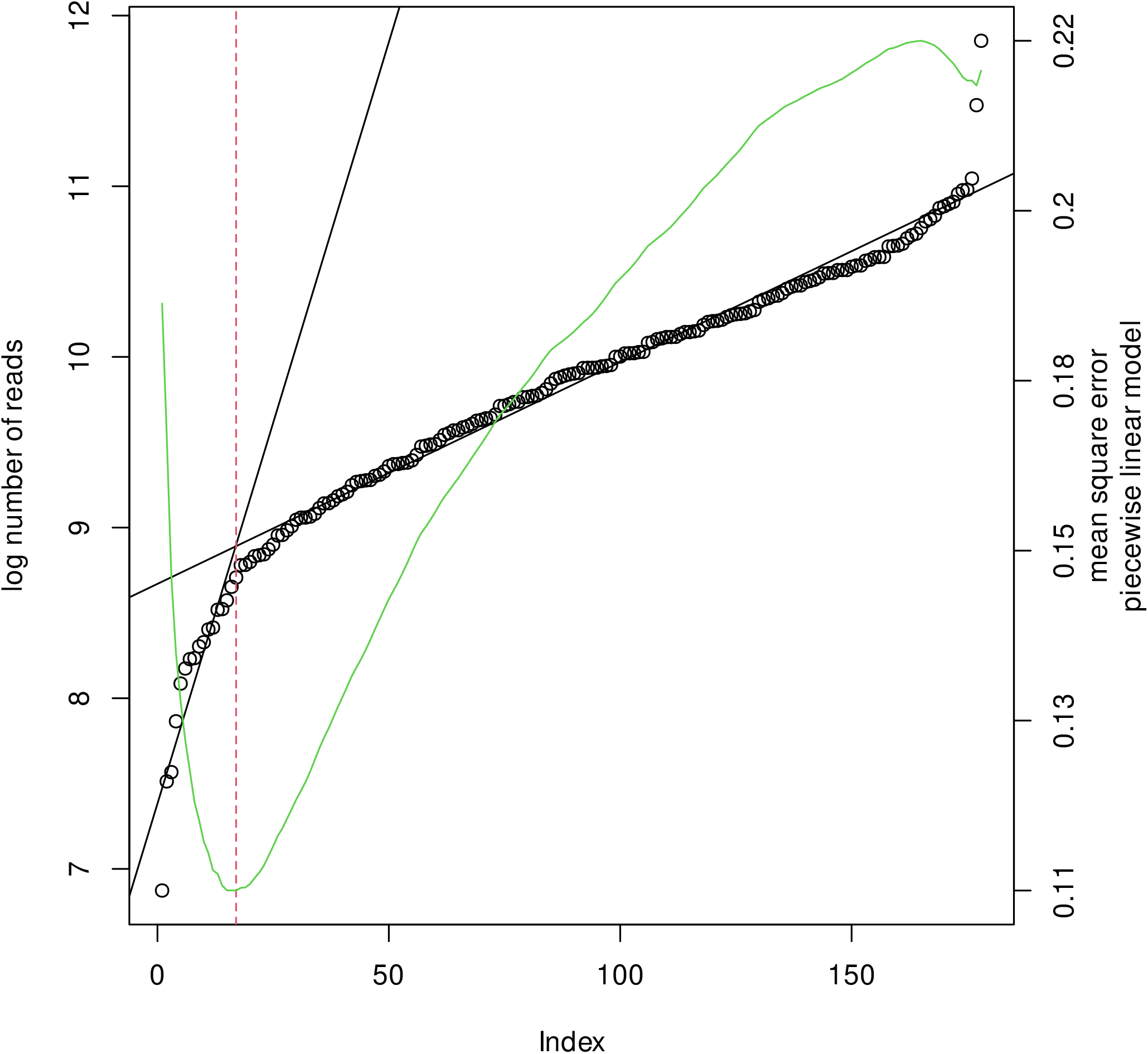
Determination of the threshold to remove samples with too few reads to be included in the analyses by piecewise linear model on the log of the number of reads sorted increasingly. The two black lines represent the two linear models at the optimal breaking point (vertical red line). The green line represents the mean square error for each breaking point. The 162 samples on the right of the breaking point were kept for further analyses.

**Figure S4:**
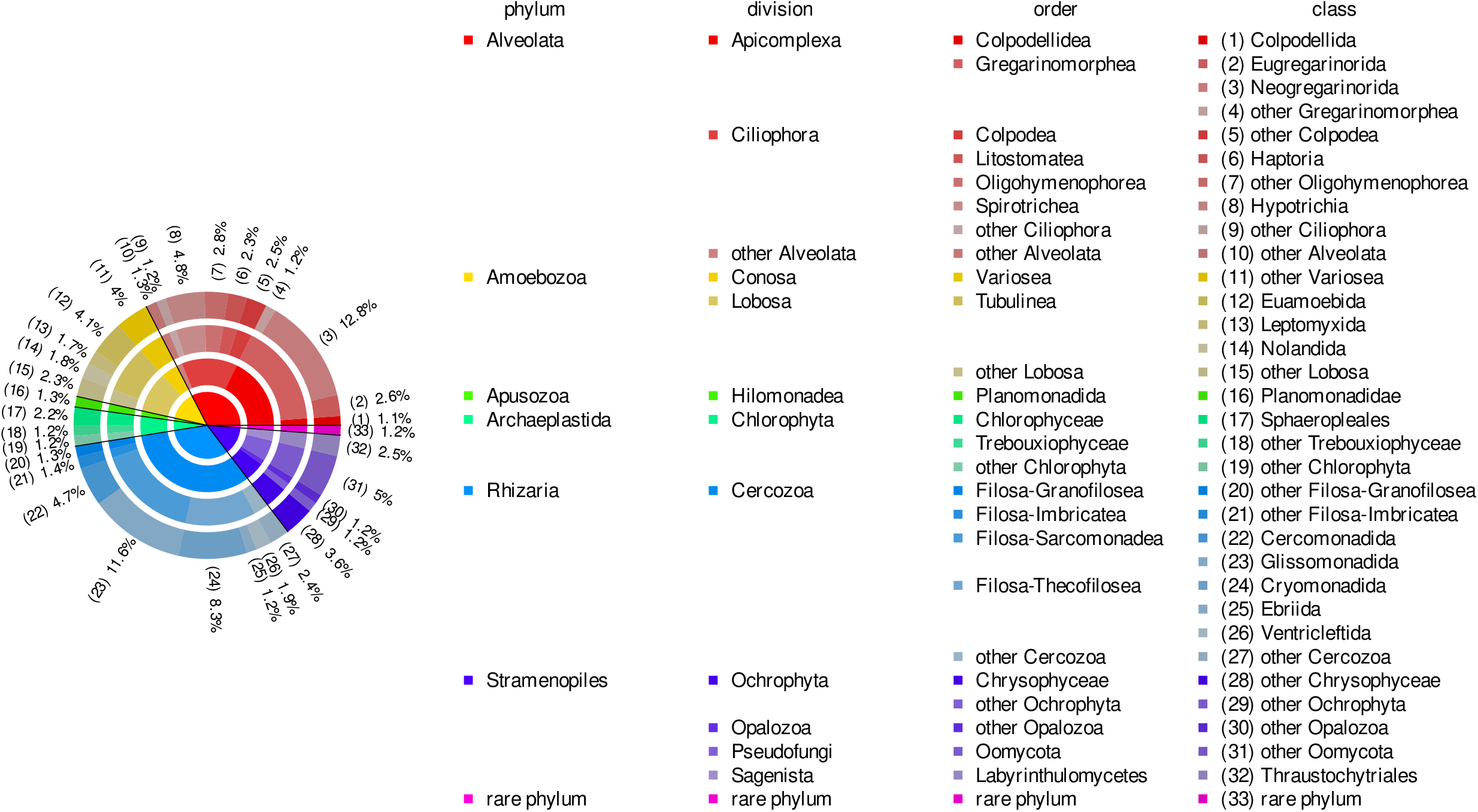
Protist taxa’s relative abundance retrieved from 162 meadow soil sampled in the Swiss western Alps (3992 OTUs, 3’334’390 sequences). The different concentric circles represent the different taxonomic levels, phylum are toward the centre whereas classes are at the exterior of the pie chart. Taxa representing less than 1% of the community were pooled within the “other” category of their lower taxonomic level.

**Figure S5:**
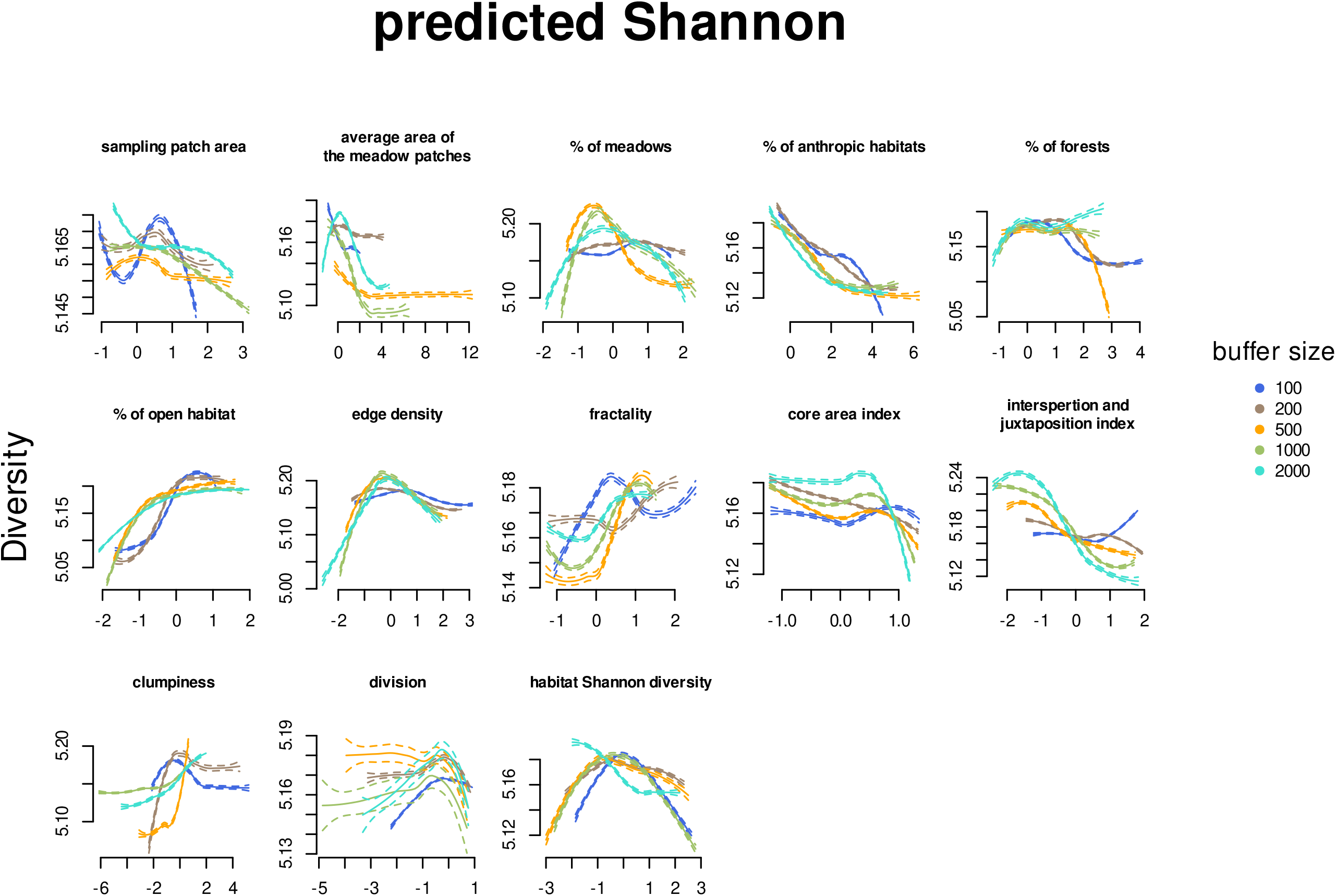
Effect of landscape descriptors on protist Shannon diversity. Partial dependence plots showing the overall protist Shannon diversity response in function of 15 landscape descriptors from meadow soils taken in the Swiss western Alps.

